# A new model of human dispersal

**DOI:** 10.1101/031674

**Authors:** Trevor G. Underwood

## Abstract

Analysis of previously unpublished allele counts obtained from the French-San-Neanderthal-Chimpanzee alignment of the high quality DNA sequence of a Neanderthal from the Altai Mountains raises significant questions about the currently accepted phylogenetic model of the origins of Europeans. Previous estimates of the proportion of Neanderthal ancestry in present-day Europeans ranged between 1.3% and 2.7% supporting a recent Out-of Africa dispersal model followed by a low level of admixture with Neanderthals. However, analysis of the allele counts indicates the existence of an unidentified third archaic ancestor of Europeans, which diverged from its common ancestor with sub-Saharan Africans around 900 thousand years ago. This analysis shows that the relative proportions of derived alleles in the 0.0826% of the European genome that is not shared with the common ancestor of humans and chimpanzee are 13.6% Neanderthal, 32.3% sub-Saharan African and 54.2% third archaic ancestor. This analysis together with anthropological and archaeological evidence suggests a new model of human dispersal based on a Eurasian lineage in the Levant, which admixed with Neanderthals and descendants of African mtDNA haplogroup L3, followed by radiation from a basal admixed population around 55–50 Kya, with no subsequent major contribution to the European genome.

## Article

This paper examines previously unpublished allele counts obtained from the French-San-Neanderthal-Chimpanzee alignment of the high quality DNA sequence of a Neanderthal from the Altai Mountains. This analysis indicates the existence of a third archaic ancestor of present-day Europeans, which diverged from its common ancestor with sub-Saharan Africans around 900 thousand years ago. It also shows that the relative proportions of derived alleles in the 0.0826% of the European genome that is not shared with the common ancestor of humans and chimpanzee are 13.6% Neanderthal, 32.3% sub-Saharan African and 54.2% third archaic ancestor. This differs significantly from the currently accepted phylogenetic model of the origins of Europeans based on a recent Out-of Africa dispersal model, followed by a low level of admixture with Neanderthals. If this new model of human dispersal is correct, it has profound implications for the interpretation of the anthropological and archaeological evidence, which has largely been framed within the Out-of-Africa model^1^.

Humans have 23 pairs of chromosomes in the nucleus of each cell; one of each pair inherited from the mother and the other from the father. Chromosomes consist of long strands of (double-stranded) nuclear DNA, comprising nitrogenous bases and a sugar-phosphate backbone. There are four possible nitrogenous bases, adenine, guanine, cytosine, and thymine, at each location (locus) on a single strand of the DNA molecule. The variants are referred to as alleles. A sequence of three nucleic acids in the DNA molecule specifies which amino acid will be created during protein synthesis. Proteins determine the phenotype, e.g. red hair. During reproduction the pairs of strands of DNA (one from the mother and the other from the father of each individual) are copied, then shuffled during the production of the egg and sperm, and recombined into two separate cells. After fertilization, the offspring inherits a unique combination of the DNA of all four grandparents. Changes in alleles, referred to as derived alleles, occur as the result of mutations, caused by copy errors in reproducing DNA, which are then passed on to the next generation. Within an archaic lineage most of the derived alleles of that lineage are shared by all grandparents. New mutations may or may not be passed on depending on the reproductive success of the offspring. DNA, known as mitochondrial DNA or mtDNA, is also present outside the nucleus, in the mitochondria. This is inherited solely from the mother.

Sequencing refers to identifying the allele at each locus on a string of DNA. Alignment involves lining up DNA sequences of different species or subspecies, such as a French, San, Neanderthal and Chimpanzee individual, to identify differences in the variant of the nitrogenous base (allele) at each locus on a chromosome. This is known as the allele pattern. An allele that is shared with the common ancestor of humans and chimpanzee is described as ancestral and labelled A, and derived alleles are labelled B, C or D, as there can be up to four different alleles corresponding to the four nitrogenous bases. The allele pattern AAAA at a locus in the French-San-Neanderthal-Chimpanzee alignment corresponds to the allele being ancestral for each population in the alignment. BABA corresponds to the French and Neanderthal sharing a derived allele, and the San and the Chimpanzee sharing the ancestral allele. The number of occurrences of each allele pattern over the whole genome (i.e. all chromosomes), for loci where the allele is identified for each genome, is known as the allele count for that allele pattern.

A major advance in understanding the origins of present-day *Homo sapiens* was made possible by the first sequencing of the nuclear genome of Neanderthals from three Neanderthal bones from Vindija Cave in Croatia by a group led by Svante Pääbo at the Max Planck Institute for Evolutionary Anthropology in Leipzig (Pääbo 2014). This was published in Green et al (2010) in *Nature* on May 7, 2010. This sequence was aligned with the sequences of the genomes of various present-day non-Africans, including a French; present-day sub-Saharan Africans who had remained isolated from non-Africans, including a San; and a chimpanzee. The Neanderthal DNA obtained from the Vindija Cave was not of very high quality, resulting in a significant number or errors when reading the Neanderthal sequence, but a much higher quality sample of DNA was obtained from a 50 Kya Neanderthal proximal pedal phalanx toe bone from Denisova Cave in the Altai Mountains in southern Siberia (correspondence with Nick Patterson, February 5, 2014). This was sequenced by the same group and the results were published in Prüfer et al (2014) in *Nature* on January 2, 2014.

In Green et al (2010) and Prüfer et al (2014), David Reich and Nick Patterson introduced the D statistic, based on the difference between the allele counts for allele patterns BABA and ABBA in the European-African-Neanderthal-Chimpanzee alignment, where the European and Neanderthal share the same derived alleles in BABA and the African and Neanderthal share the same derived alleles in ABBA, to determine whether Neanderthals were more closely related to non-Africans than to Africans (Green et al 2010, Suppl. 15; Prüfer et al 2014, Suppl. 14). See Appendix A.

The allele counts for the BABA and ABBA allele patterns were then used to estimate the admixture proportion using two different estimators obtained by substituting statistics based on these allele patterns in the two population admixture equation *E* = *fN_A_* + *(1−f) A_A_,* where *E* is the present-day European genome, *N_A_* is the Neanderthal genome that contributed alleles to the present-day European, *A_A_* is the sub-Saharan African ancestral genome that contributed alleles to the present-day European, and *f* is the Neanderthal admixture proportion (Green et al 2010, Suppl. 15, equation S15.7; Suppl. 18, equation S18.1; Prüfer et al 2014, Suppl. 14, equation S14.8).

In Green et al (2010), the S statistic, equal to the numerator of the D statistic, was substituted for each population in the two population admixture equation to derive an admixture estimator, *f* as a ratio of two S statistics (Appendix A). This was interpreted as measuring the relative rate of matching of the European and African sequences to the Neanderthal sequence. In Prüfer et al (2014), the F_4_ statistic, seen to measure the overlap between the drift paths of two admixing populations, was substituted for each population in the two population admixture equation to derive the *f*_4_-ratio admixture estimator as a ratio of the F_4_ statistics on each side of the phylogeny (Appendix A).

In Green et al (2010), Reich and Patterson estimated that the proportion of Neanderthal ancestry in present-day Europeans ranged between 1.3% and 2.7% (Green et al. 2010, Suppl. 18). In Prüfer et al (2014), their estimates ranged between 1.48% and 1.96% (Prufer et al. 2014, Suppl. 14). Neanderthal admixture was defined as “the proportion of lines of descent [or] probability that the line of descent, will pass through the Neanderthal population [or] lineage”, or “the proportion that traces its genealogy through the Neanderthal side of the phylogenetic tree” (correspondence with Nick Patterson, June 17, 2012, and January 24, 2014; Green et al. 2010, Suppl. 18).

Appendix A examines how the substitution of the S statistic and the F_4_ statistic in the two population admixture equation resulted in estimators which are not equal to the admixture proportion, and how the estimators actually used by Reich and Patterson to calculate the admixture proportion do not correspond to their definitions of Neanderthal admixture (Appendix A. Mathematical error in the derivation of estimators of admixture proportions).

### Allele counts

This paper takes a different approach by first examining the allele counts obtained from the French-San-Neanderthal-Chimpanzee alignments to see what they might tell us about the contributions of archaic ancestors to the European genome. The allele counts for the French-San-Vindija Neanderthal-Chimpanzee alignment were published in Green et al (2010), Suppl. 15, Table S51. A similar set of allele counts for the French-San-Altai Neanderthal-Chimpanzee alignment were kindly provided to the author by Nick Patterson after the publication of the sequencing of the high quality DNA from the Altai Neanderthal in Prüfer et al (2014). See Table 1. The analysis in this paper focuses on the previously unpublished allele counts from Prüfer et al (2014), but also references the allele counts from the Vindija alignment to demonstrate consistency and variation in sequencing errors between the two alignments.

**Table 1.** Counts of all possible allele patterns in the two alignments

| Allele pattern | Relabeled # | French-San-Vindija Neanderthal-Chimpanzee allele counts* | French-San-Altai Neanderthal-Chimpanzee allele counts** |
| --- | --- | --- | --- |
| AAAA | AAAA | 818,322,920 | 1,389,787,867 |
| AABA | AABA | 5,827,247 | 755,369 |
| ABAA | ACAA | 689,594 | 647,939 |
| ABBA | ABBA | 95,347 | 143,385 |
| ABCA | ACAA## | 2,995 | 465 |
| BAAA | DAAA | 756,324 | 629,542 |
| BABA | BABA | 103,612 | 157,935 |
| BACA | DAAA## | 3,544 | 386 |
| BBAA | CCAA | 303,340 | 375,289 |
| BBBA | AAAE | 8,156,936 | 16,332,396 |
| BBCA | AAAE## | 32,607 | 11,376 |
| BCAA | DAAA## | 972 | 405 |
| BCBA | AAAE## | 6,147 | 8,237 |
| CBBA | AAAE## | 6,264 | 7,693 |
| BCDA | DAAA## | 36 | 10 |
| Total |  | 834,307,885 | 1,408,858,294 |

These allele counts offer the possibility of an extraordinary insight into the genetic makeup and ancestral history of present-day Europeans. The fifteen allele counts, covering all possible allele patterns from the Altai alignment, were obtained from more than one billion data quality filtered sites on the respective genomes of the four populations, so potentially provide an extremely reliable source of information. In spite of real possibilities of sequencing errors, particularly with the poorer quality archaic DNA samples, it seems reasonable to start by assuming that the allele counts are substantially correct, although this assumption will be re-examined later.

In order to analyze this data it is helpful to understand what these allele patterns represent and relabel them accordingly so the letters correspond to the most likely source of the allele in the European genome. For this purpose, we will designate A = ancestral to humans and chimpanzee, B = Neanderthal, C = sub-Saharan African, D = European, and E = Chimpanzee differs from common ancestor of humans. We can then try to deduce what the observed counts tell us.

For example, the allele pattern BABA indicates that at these sites the San and Chimpanzee share the ancestral allele A and the French and Neanderthal share a derived allele B, which we will assume to have originated in an ancestor of the Neanderthal, although as we will see later it could have originated in another archaic ancestor of the French which admixed with an ancestor of the Neanderthal. Consequently, this will retain its label BABA. However, the allele pattern labelled BBBA relative to the Chimpanzee being considered ancestral, which includes both mutations AAAB that arose in the Chimpanzee genome since the split with humans up until the present time and mutations BBBA that arose in the common ancestor of humans after the split with Chimpanzee but before the divergence of archaic human ancestors, will be relabeled AAAE to include these with other ancestral alleles of humans in the European genome. The amended labels are shown in the second column of Table 1.

### Error in the Vindija AABA allele count

The first point to note is that comparison of the alignments using the Croatian Vindija and Siberian Altai Neanderthals indicates a significant similarity in allele patterns obtained using these geographically distant Neanderthal DNA samples, providing confidence about the validity of the counts. The main exception is the allele count for AABA, which is 5,827,247 (0.7% of the total) in the Vindija alignment versus 755,369 (0.05% of the total) in the Altai alignment (Table 1). This allele pattern represents mutations in the Neanderthal genome at sites where the French, San and Chimpanzee alleles are the same as each other, and therefore presumably ancestral. Because the mutation in this allele pattern derives from the sequencing of the archaic Neanderthal DNA, this is particularly vulnerable to sequencing error, given that a misread of the Neanderthal allele as B rather than A is more likely than any other error due to the very large AAAA count. It is also more likely that the error will be larger in the sequencing of the low quality Vindija DNA samples compared with the high quality Altai DNA.

This is confirmed by comparing these AABA counts with the corresponding relabeled ACAA counts, which represent mutations in the San genome at sites where the French, Neanderthal and Chimpanzee are the same as each other, and therefore presumably ancestral. In a population model where Neanderthals and sub-Saharan Africans share a common ancestor, we would expect the allele counts AABA and ABAA to be approximately equal within each alignment as these counts indicate the divergence time of the two populations. In the Vindija alignment, the AABA count of 5,827,247 is significantly different from the ABAA count of 689,594; whereas in the Altai alignment these counts are similar; 755,369 and 648,304, respectively. This suggests a very large sequencing error in Vindija AABA count; and a considerably reduced error in this most vulnerable allele pattern in the Altai sample. This is very reassuring with regard to our assumption that the Altai alignment allele patterns are substantially correct. Consequently, unless specifically identified, the rest of this analysis will refer to the allele counts from the Altai alignment.

### Allele pattern AAAE; percentage of human genome shared with the chimpanzee

For the analysis of the European genome, the next step is to reorder the allele patterns so they are grouped according to whether the French (European) allele is ancestral A, or the same as the derived Neanderthal allele B, or the same as the derived San (sub-Saharan African) allele C, or represents a unique derived European allele D (Table 2).

**Table 2.** Analysis of allele patterns/adjusted branch lengths: French-San-Altai Neanderthal-Chimpanzee

| Allele pattern | Allele counts | % | Divergence time/branch length<br>using human mutation rate<br>(bp/year):<br>(.5 x 10<sup>-9</sup>) (1.0 x 10<sup>-9</sup>) |  |  | Mutation rate<br>calculated from<br>allele count ##<br>(bp/year) | Sequence<br>error rate<br>(%) | Adjusted<br>branch<br>length ***<br>(.5 x 10<sup>-9</sup>) |
| --- | --- | --- | --- | --- | --- | --- | --- | --- |
| Ancestral alleles of humans in European genome: |  |  |  |  |  |  |  |  |
| AAAA | 1,389,787,867 | 98.6464 |  |  |  |  |  |  |
| AAAE (+ BBBA) | 16,332,396 | 1.1593 | 12,478,691 | 6,239,345 | ** | .000000000500 |  |  |
| BCBA = AAAE # | 8,237 | .0006 |  |  | C* | .000000000540 | 7.9 |  |
| BBCA = AAAE # | 11,376 | .0008 |  |  | B* | .000000000612 | 22.3 |  |
| CBBA = AAAE # | 7,693 | .0005 |  |  | D* | .000000000504 | .8 |  |
| AABA | 755,369 | .0536 | 1,087,028 | 543,514 | * | .000000000500 |  | 888,725 |
| ABAA = ACAA | 647,939 | .0460 | 932,954 | 466,477 | * | .000000000500 |  | 863,877 |
| ABCA = ACAA# | 365 | .0000 |  |  | B* | .000000000603 |  |  |
| ABBA | 143,385 | .0102 | 206,341 | 103,170 | * |  |  |  |
| A*** | 1,407,694,627 | 99.9174 |  |  |  |  |  |  |
| A*** – AAAE | 1,391,334,925 | 98.7562 |  |  |  |  |  |  |
|  |  | (98.77-98.94%) | Mikkelsen et al (2005) |  |  |  |  |  |
| Neanderthal ancestor derived alleles in European genome: |  |  |  |  |  |  |  |  |
| BABA | 157,935 | .0112 |  |  |  |  |  |  |
| B*BA | 157,935 | .0112 | 227,279 | 113,640 | * |  |  |  |
| (B*BA)*100/(B*BA+CC*A+D**A) |  | 13.6 |  |  |  |  |  |  |
| African ancestor derived alleles in European genome: |  |  |  |  |  |  |  |  |
| BBAA = CCAA | 375,289 | .0266 |  |  |  |  |  |  |
| CC*A | 375,289 | .0266 | 540,067 | 270,033 | * |  |  |  |
| (CC*A)*100/(B*BA+CC*A+D**A) |  | 32.3 |  |  |  |  |  |  |
| Other derived alleles in European genome: |  |  |  |  |  |  |  |  |
| BAAA = DAAA | 629,542 | .0447 |  |  |  |  |  |  |
| BACA = DAAA# | 386 | .0000 |  |  | B* | .000000000656 |  |  |
| BCAA = DAAA# | 405 | .0000 |  |  | C* | .000000000689 |  |  |
| BCDA = DAAA# | 10 | .0000 |  |  | BC* | .000000000686 |  |  |
| D**A | 630,343 | .0447 | 907,107 | 453,553 | * | .000000000500 |  | 899,846 |
| D**A*100/(B*BA+CC*A+D**A) |  | 54.2 |  |  |  |  |  |  |
| Total alleles | 1,408,858,194 |  |  |  |  |  |  |  |

The first observation from this table is that the total number of alleles for which the French allele is ancestral (relabeled allele patterns A***) indicates that 99.9174% of the French (European) genome is ancestral. Subtracting the allele count for AAAE, where the common ancestor of humans and Chimpanzee differ, results in 98.7562% of the European genome being identical to that of the Chimpanzee. This corresponds very closely with the range 98.77–98.94% reported by the Chimpanzee Consortium in Mikkelsen et al (2005), so again this is very reassuring. Assuming the currently most favored human mutation rate of 0.5 × 10^−9^ bp per year^2^, the AAAE allele count corresponds to a genetic divergence time between humans and chimpanzee of ((16,332,396/1,389,787,867)/0.5 × 10^−9^ + 1,400,000 + 50,000)/2 = 12.5 Mya, allowing 1.4 My for the branch shortening arising from the subsequent split between ancestors of Neanderthals, sub-Saharan Africans and Europeans and 50 Ky for the date of the Neanderthal sample (see Genetic Divergence Times below and Table 2). A human mutation rate of 1.0 × 10^−9^ bp per year would halve this estimate to approximately 6.2 Mya.

### Allele patterns AABA, ACAA and DAAA: existence of third archaic ancestor of Europeans

On examining the remaining allele patterns, the first major question concerns the allele pattern DAAA, for which the French allele is different from the San, Neanderthal and Chimpanzee alleles and for which the allele count is 630,343, about the same as AABA at 755,369 and ACAA at 648,304 and. As noted previously, AABA represents mutations in the Neanderthal genome at sites where the French, San and Chimpanzee alleles are the same as each other, and therefore presumably ancestral, and ACAA represent mutations in the San genome at sites where the French, Neanderthal and Chimpanzee are the same as each other, and therefore presumably ancestral. Similarly, DAAA represents mutations in the French genome at sites where the San, Neanderthal and Chimpanzee alleles are the same as each other, and therefore presumably ancestral; but what does this mean?

The two population admixture equation which was used in the derivations of the admixture estimators assumed that the present-day European genome is an admixture of a Neanderthal-related genome and a sub-Saharan African-related genome, *E* = *fN_A_* + *(1−f) A_A_.* In this model, mutations in the European genome are either mutations in the ancestral Neanderthal genome or mutations in the ancestral sub-Saharan African genome and cannot contribute to DAAA as they would also be present in Neanderthals or sub-Saharan Africans. So the DAAA allele pattern can only represent mutations in the European genome (i.e. A to D in AAAA resulting in DAAA) occurring after the admixture event. However, the number of such mutations far exceeds what is possible under the usual assumption of an approximately constant rate of human mutation and admixture between Neanderthals and African ancestors being relatively recent (within the last two hundred thousand years).

The AABA count corresponds to a branch length on the Neanderthal lineage of (755,369/1,389,787,867)/(0.5 × 10^−9^) = 1,087 Ky, assuming a human mutation rate of 0.5 × 10^−9^ bp per year; or 1,137 Ky, allowing for the Altai Neanderthal sample being 50 Kya old. The ACAA count corresponds to a branch length on the African lineage of (648,304/1,389,787,867)/(0.5 × 10^−9^) = 933 Ky; and the DAAA allele count corresponds to a branch length on the European lineage of (630,343/1,389,787,867)/(0.5 × 10^−9^) = 907 Ky (Table 2)^3^.

### Adjustment for sequencing errors

Allele patterns, such as BCBA, with more than two alleles could either reflect sequencing errors or two or more separate mutations at the same single nucleotide polymorphism (SNP). In tables 2, 3 and 4, these were adjusted by replacing the allele that is most likely in error or resulted from a second or third mutation with the allele of most likely allele pattern, and relabeling accordingly. For example, BCBA is most likely to be a misread or mutation of BBBA, which was relabeled as AAAE.

**Table 3.** Analysis of allele patterns: French/Ust’Ishim/Kostenki-San-Altai Neanderthal-Chimpanzee

| Table 3 |  |  |  |  |  |  |
| --- | --- | --- | --- | --- | --- | --- |
| Allele pattern | Allele counts<br>French* | % | Allele counts<br>Ust'-Ishim** | % | Allele counts<br>Kostenki 14*** | % |
| Common ancestral alleles: |  |  |  |  |  |  |
| AAAA | 1,389,787,867 | 98.6464 | 1,028,702 | 42.4708 | 7,163,742 | 59.4574 |
| BBBA = AAAE | 16,332,396 | 1.1593 | 1,034,962 | 42.7293 | 652,675 | 5.4170 |
| BCBA = AAAE# | 8,237 | 0.0006 |  | 0.0000 |  | 0.0000 |
| BBCA = AAAE# | 11,376 | 0.0008 |  | 0.0000 |  | 0.0000 |
| CBBA = AAAE# | 7,693 | 0.0005 |  | 0.0000 |  | 0.0000 |
| AABA | 755,369 | 0.0536 | 102,587 | 4.2354 | 1,234,858 | 10.2490 |
| ABAA = ACAA | 647,939 | 0.0460 | 86,155 | 3.5570 | 1,048,857 | 8.7053 |
| ABCA = ACAA# | 365 | 0.0000 |  | 0.0000 |  | 0.0000 |
| ABBA | 143,385 | 0.0102 | 18,866 | 0.7789 | 235,769 | 1.9568 |
| A*** | 1,407,694,627 | 99.9174 | 2,271,272 | 93.7714 | 10,335,901 | 85.7856 |
| A*** - AAAE | 1,391,334,925 | 98.7562 | 1,236,310 | 51.0421 | 9,683,226 | 80.3685 |
| *Mikkelsen et al (2005) (98.77-98.94%)* |  |  |  |  |  |  |
| Neanderthal ancestor derived alleles: |  |  |  |  |  |  |
| BABA | 157,935 | 0.0112 | 21,536 | 0.8891 | 277,136 | 2.3002 |
| B*BA | 157,935 | 0.0112 | 21,536 | 0.8891 | 277,136 | 2.3002 |
| (B*BA)*100/(B*BA+CC*A) |  | 29.6 % |  | 31.0 % |  | 32.9 % |
| (B*BA)*100/(B*BA+CC*A+D**A) |  | 13.6 % |  | 14.3 % |  | 16.2 % |
| African ancestor derived alleles: |  |  |  |  |  |  |
| BBAA = CCAA | 375,289 | 0.0266 | 47,852 | 1.9756 | 566,140 | 4.6988 |
| CC*A | 375,289 | 0.0266 | 47,852 | 1.9756 | 566,140 | 4.6988 |
| (CC*A)*100/(B*BA+CC*A) |  | 70.4 % |  | 69.0 % |  | 69.0 % |
| (CC*A)*100/(B*BA+CC*A+D**A) |  | 32.3 % |  | 31.7 % |  | 33.1 % |
| Other derived alleles: |  |  |  |  |  |  |
| BAAA = DAAA | 629,542 | 0.0447 | 81,478 | 3.3639 | 869,356 | 7.2155 |
| BACA = DAAA# | 386 | 0.0000 |  | 0.0000 | 0 | 0.0000 |
| BCAA = DAAA# | 405 | 0.0000 |  | 0.0000 | 0 | 0.0000 |
| BCDA = DAAA# | 10 | 0.0000 |  | 0.0000 | 0 | 0.0000 |
| D**A | 630,343 | 0.0447 | 81,478 | 3.3639 | 869,356 | 7.2155 |
| D**A*100/(B*BA+CC*A+D**A) |  | 54.2 % |  | 54.0 % |  | 50.8 % |
| Total alleles | 1,408,858,194 |  | 2,422,138 |  | 12,048,533 |  |
| # More than two alleles; substituted with most likely allele pattern |  |  |  |  |  |  |
| Relative branch lengths: |  |  |  |  |  |  |
| AABA | 755,369 | 37.2 | 102,587 | 38.0 | 1,234,858 | 39.2 |
| ABAA = ACAA | 647,939 | 31.9 | 86,155 | 31.9 | 1,048,857 | 33.3 |
| BAAA = DAAA | 629,542 | 31.0 | 81,478 | 30.2 | 869,356 | 27.6 |
|  | 2,032,850 |  | 270,220 |  | 3,153,071 |  |
| Divergence times (years) = adjusted branch lengths + time of death (human mutation rate (base pair/yr) = $0.5 \times 10^{-9}$). | | | | | | |
| Neanderthal (AABA + BABA + ABBA + 50 Ky) | 1,372,345 |  |  |  |  |  |
| San (ACAA + CCAA) | 1,404,469 |  | San (ACAA) | 864,402 |  |  |
| French (DAAA + CCAA) | 1,439,913 |  | French (DAAA) | 899,846 |  |  |
| * Data provided by Nick Patterson from the alignments in Prüfer et al (2014). |  |  |  |  |  |  |
| ** Data provided by Qiaomei Fu from the alignments in Fu et al (2014). |  |  |  |  |  |  |
| *** Data provided by Martin Sikora from the alignments in Seguin-Orlando et al (2014). |  |  |  |  |  |  |

**Table 4.**
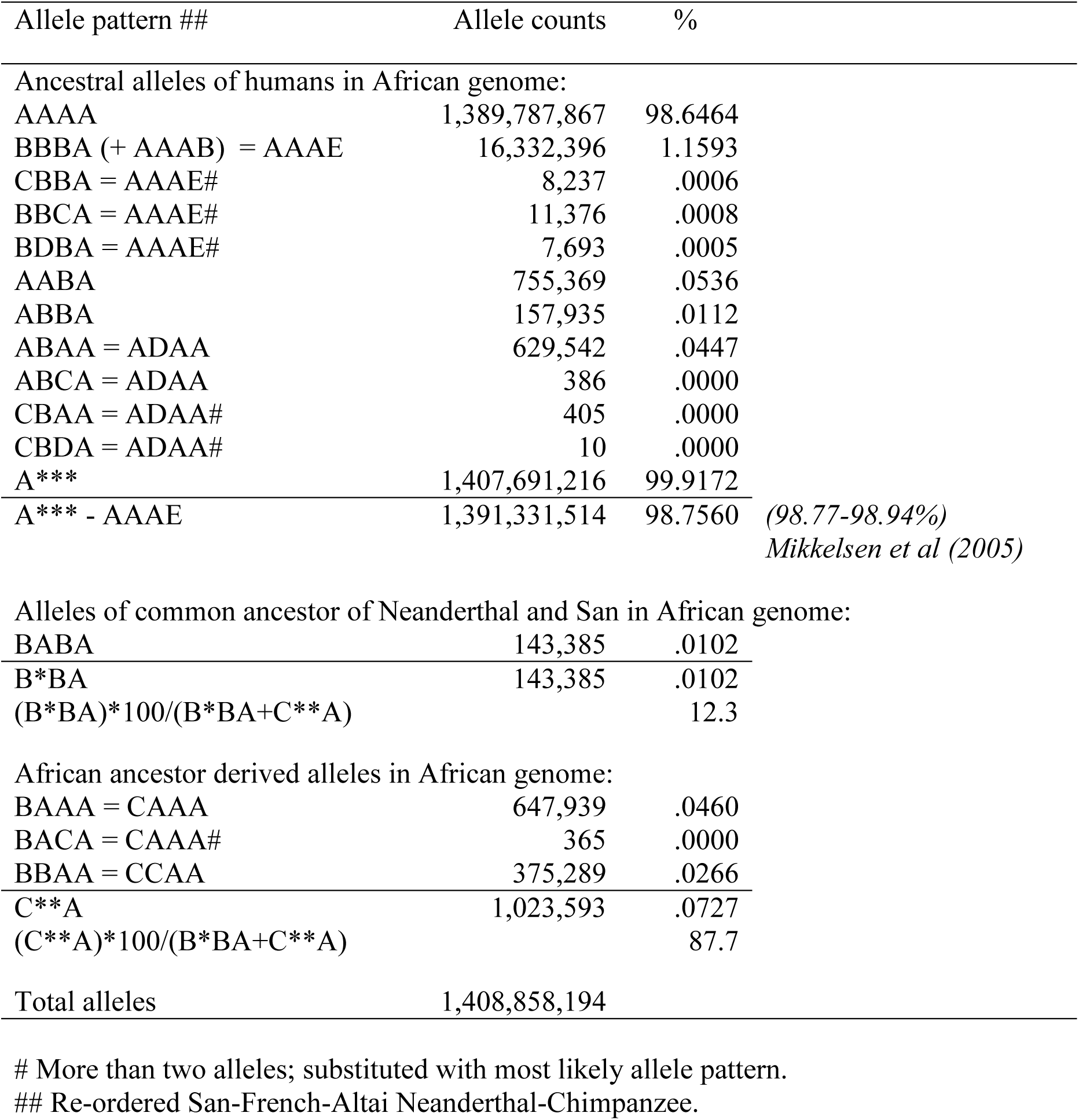
Analysis of allele patterns: San-French-Altai Neanderthal-Chimpanzee

However, calculation of the mutation rate required to create these allele counts suggests that they are largely consistent with two separate mutations, or in the case of BCDA, three separate mutations, with a relatively small contribution from sequencing errors. As might be expected, the implied residual sequencing errors are largest in the Altai Neanderthal sample (22.3% of the mutation rate), less in the San (7.9%), and least in the French (0.8%) (Table 2).

Adjusting the previously calculated branch lengths for the three lineages for the corresponding implied sequencing error rates above results in branch lengths of around 889 Ky for the ancestor of Neanderthals, corresponding to AABA, or 939 Ky allowing for the age of the Altai Neanderthal sample; of around 864 Ky for the ancestor of present-day sub-Saharan Africans, corresponding to ACAA; and of around 900 Ky for the unidentified third ancestor of present-day Europeans, corresponding to DAAA (Table 2). As AABA, ACAA and DAAA represent mutations unique to the respective populations, these adjusted branch lengths correspond to the time during which the ancestral populations of Neanderthals, Africans and Europeans were distinct from each other. The fact that they are approximately equal provides further support for this analysis and the data.

Consequently, the most parsimonious explanation of the DAAA count is the existence of a previously unidentified third archaic ancestor of Europeans. This implies a revised three population admixture equation *E* = *f^N^N_A_* + *f^A^A_A_* + *(1−f^N^ − f^A^)O_A_,* where *O_A_* is the genome of the third archaic ancestor of Europeans. This further invalidates the derivation of the admixture estimators used in Green et al (2010) and Prüfer et al (2014), which assumed a two population admixture equation *E* = *fN_A_* + *(1−f)A_A_*.

### Allele patterns CCAA, BABA and ABBA; shared mutations and admixture

The CCAA count of 375,289, indicating (375,289/1,389,787,867)/(0.5 × 10^−9^) = 540 Ky of mutations, represents either shared mutations after the common ancestor of Europeans and Africans had split from the ancestor of Neanderthals and before their ancestors split from each other, or the sharing of mutations on the European or African lineages from more recent admixture between the ancestors of Europeans and Africans (Table 2.) This allele count represents an additional 540 Ky of mutations on the European and/or African lineages.

Under the two archaic ancestor model, all mutations on the shared European and African lineage up until their split around 50 Kya should result in CCAA allele counts, and this should be much larger than either the DAAA or ACAA allele counts, representing mutations since the split between Europeans and Africans. The fact that the DAAA and ACAA allele counts, respectively 629,542 and 647,939 and representing around 900 Ky of mutations, both of which should be free of significant sequencing errors, are approximately equal in size and significantly larger than the CCAA allele count provides further evidence of the third archaic ancestor of Europeans and that the two archaic ancestor model for Europeans is incorrect.

The BABA allele count, with the same derived allele at a locus in the European and Neanderthal genomes, and African and Chimpanzee sharing the ancestral allele, of 157,935, indicating (157,935/1,389,787,867)/(0.5 × 10^−9^) = 227 Ky of mutations, most likely reflects the remnants of gene flow from Neanderthals into the third archaic European ancestor between 250–55 Kya^4^.

The ABBA allele count, with the same derived allele at a locus in the African and Neanderthal genomes, and European and Chimpanzee sharing the ancestral allele, of 143,385, indicating (143,385/1,389,787,867)/(0.5 × 10^−9^) = 206 Ky of mutations, probably reflects the remnants of gene flow from a Neanderthal ancestor, probably *Homo heidelbergensis,* into the African ancestor around 600 Kya (Table 2). There is evidence that a group similar to European *Homo heidelbergensis*, sometimes referred to as *Homo rhodesiensis* to distinguish it from the European variety, was present in Africa between 600–200 Kya (Rightmire 1996)^5^. It may be that *Homo heidelbergensis* expanded into Africa and admixed with the archaic African ancestor.

The BABA and ABBA allele counts contribute an additional 227 Ky + 206 Ky = 433 Ky to the divergence time on the Neanderthal lineage. Under the two archaic ancestor model, the BABA and ABBA allele counts can only be explained by polymorphisms, incomplete lineage sorting or linkage disequilibrium (see Alternative Explanations below.)

### Genetic divergence times and the archaeological and anthropological evidence

Allowing for time of death of the Neanderthal of around 50 Kya, and assuming that the BABA and ABBA allele patterns represent gene flow from Neanderthals into the third archaic ancestor, and gene flow from Neanderthal ancestors into African ancestors, respectively; and the CCAA allele patterns represent shared mutations after the genetic divergence of the common ancestor of Europeans and Africans from the ancestor of Neanderthals and before the split between the African ancestor and the third archaic ancestor of Europeans, this would result in genetic divergence times for the three lineages as follows (Table 3): Neanderthal = AABA + BABA + ABBA + 50 = 889 Ky + 227 Ky + 206 Ky + 50 Ky = 1,372 Kya; African = ACAA + CCAA = 864 Ky + 540 Ky = 1,404 Ky; and European third archaic ancestor = DAAA + CCAA = 900 Ky + 540 Ky = 1,440 Kya.

This would also indicate an initial split between the common ancestor of Europeans and Africans and the ancestor of Neanderthals around 1.4 Mya, followed by a split between the ancestor of Africans and the third archaic ancestor of Europeans around 900 Kya^6^. These genetic divergence times, which are, of course, subject to the accuracy of the estimated human mutation rate, correspond very well with the archaeological and anthropological evidence, providing further support for this analysis and for the average mutation rate.

Although there is evidence of *Homo erectus* or other hominins in Africa, Asia, and Europe between 2 Mya and 1.4 Mya, the DNA divergence dates suggest that the archaic ancestors of Neanderthals, sub-Saharan Africans and Europeans derived from a common *Homo erectus* ancestor around 1.4 Mya, probably a group of sub-Saharan African ancestors with Oldowan I technology expanding out of Africa and continuing northward and westward into Europe^7^.

The genetic divergence dates indicate that descendants of this population subsequently expanded out of Africa into the Levant around 900 Kya, forming the third archaic lineage^8^. Although the common ancestor could have been *Homo erectus* in Eurasia with back migration into Africa, given the widespread presence of Acheulian artifacts and fossil evidence of *Homo erectus (ergaster)* in Africa between 1.4 Mya and 900 Kya, the re-emergence of a group of *Homo erectus* from Africa around 900 Kya is the most plausible explanation.

The genetic evidence indicates that after these divergences the three lineages remained largely distinct until they coalesced in the Levant to form early Eurasian *Homo sapiens* and the Neanderthal lineage became extinct. Archaeological, anthropological and mtDNA evidence suggests that this coalescence occurred in the Levant between 250–50 Kya, and that demise of the Neanderthal lineage occurred around 40 Kya (Higham et al 2014).

### Alternative explanations

This is a fairly radical conclusion so consideration of other possible explanations is in order. Reviewers’ comments and correspondence with the authors of the admixture estimates have focused on errors in the data or on the interpretation of the data.

It has been argued that the critical DAAA allele count and the other singletons, the ACAA and AABA allele counts, are very sensitive to sequencing error because of the large number of AAAA sites which can result in these errors by a single misread compared with the ABBA and BABA counts used in the estimations based on a ratios of two S statistics or the *f*_4_-ratio. Although the AABA allele count, depending on the accurate reading of alleles in ancient DNA, is highly vulnerable, as was identified above for the low quality Vindija Neanderthal DNA, this is less true for the high quality Altai Neanderthal, and much less true of the DAAA and ACAA allele counts. The likelihood of sequencing error is least of all for the DAAA allele count, which is based on sequencing present-day European DNA, which is of far greater quality and availability than archaic DNA. As noted previously, Green et al (2010), Prufer et al (2014), and the analysis of triallelic sites above, all demonstrated that sequencing errors were low in the DAAA and ACAA allele counts.

A second argument is that the DAAA count could be the result of ancient polymorphisms in present-day human DNA. A polymorphism occurs where there is more than one variant of the nucleotide at a locus, known as an allele. Although mutations will generally result in polymorphisms when they arise (because the mutation initially affects only one individual in the population and only one of the base pairs in the double stranded DNA), over time most polymorphisms are believed to lose one or the other allele due to genetic drift or selection and become fixed.

It is argued that the excess DAAA allele counts represent derived alleles associated with common polymorphisms existing in present-day humans prior to the split between Europeans and Africans (around 50 Kya), where the European (French) individual selected the African derived allele and the African (San) individual selected the ancestral allele. (Although this could arise from polymorphisms in the Neanderthal ancestor this would be much more limited assuming only 2% of Neanderthal admixture.) This implies that the allele count should be labelled CAAA rather than DAAA under the convention adopted in this paper. In addition, a corresponding number of derived alleles associated with these polymorphisms would have created ACAA allele counts, where the European (French) individual selected the ancestral allele and the African (San) individual selected the derived allele. Where the derived alleles were selected by both individuals they would result in CCAA allele patterns.

According to this explanation, these polymorphisms would have been the result of mutations in the African ancestor existing prior to the split between the European and African ancestors which and had not yet become fixed. However, under the two archaic ancestor model, mutations on the human lineage during the 850 Ky following the split between the human and Neanderthals ancestors and prior to the split between Europeans and Africans are shared and would normally be expected to contribute to the CCAA allele count. It would also require a very large number of polymorphisms in the African ancestor at the time when the African and European populations diverged to generate a CAAA allele count representing 900 Ky of mutations when the population sizes were relatively small and consequently the number of polymorphisms relatively small. Consequently, the CCAA allele count would be expected to be significantly larger than the DAAA and ACAA allele counts. In the Altai Neanderthal alignment, the CCAA allele count was 375,289, DAAA was 630,343, and ACAA was 647,939, which effectively disposes of this explanation.^9^

A third argument is that the DAAA count could be the result of incomplete lineage sorting. Incomplete lineage sorting occurs when the common ancestor is polymorphic prior to its segregation into two lineages and both alleles are retained in the two lineages. If one of the daughter lineages divides again relatively soon then all three lineages may carry both alleles. Over time, each lineage will lose one or the other allele due to genetic drift or selection and, depending on which allele is retained, the resulting genomic segment may or may not match the overall lineage-level phylogenetic tree.

Under the two archaic population model, after the Neanderthal and African lineages divided, incomplete lineage sorting would occur at sites where the common ancestor was polymorphic and lineage 1 (European) retained the derived allele C, lineage 2 (African) retained the ancestral allele A and lineage 3 (Neanderthal) retained the ancestral allele A. Then the “gene tree” would not match the “species tree” for this gene segment. In this example the CAAA allele pattern does not match the species tree between the common ancestor of the European and African individuals and the ancestor of the Neanderthal.

The problem with this explanation, which is similar to the problem with polymorphisms existing prior to the split between Europeans and Africans, is the size of the DAAA allele count. It would require a very large number of polymorphisms in the common ancestor at the time of the split between the ancestors of Neanderthals and ancestors of Africans when the population sizes were relatively small. Not only is the DAAA allele count nearly twice the size of the CCAA allele count, but it is also approximately equal to the ACAA allele count, representing around 900 Ky of mutations on the African lineage.

In Green et al (2010), Reich and Patterson used the fact that a difference between the number of BABA and ABBA allele patterns cannot be explained by incomplete lineage sorting to detect gene flow. However, this did nothing to explain the size of these allele counts or the DAAA allele count. Green et al (2010) concluded that “Although gene flow from Neanderthals into modern humans when they first left sub-Saharan Africa seems to be the most parsimonious model compatible with the current data, other scenarios are also possible … we cannot rule out a scenario in which the ancestral population of present-day non-Africans was more closely related to Neanderthals than the present-day Africans due to ancient substructure within Africa” (Green et al 2010, page 722). This limitation was addressed by Yang et al (2012), who used simulations to show that the observed shapes of the site frequency spectrum, conditioned on a derived Vindija Neanderthal and an ancestral Yoruba nucleotide (doubly conditioned site frequency spectrum or dcfs), could not be explained by the ancient structure model but the recent admixture model provided a good fit for the observed dcfs for non-Africans, supporting the recent admixture hypothesis (Yang et al 2012, pages 2987–2993).

Lohse and Frantz (2014) demonstrated that by dividing the genome into short blocks and computing maximum likelihood estimates of parameters under models of admixture and ancestral population structure, they were able to conclusively reject ancestral structure in Africa. This analysis provided strong support for between 3.4% – 7.9% admixture from Neanderthals into Eurasian populations, noting that the D-statistic estimate was a lower bound whereas maximum likelihood estimates are unbiased (Lohse and Frantz 2014, pages 1241–1249)^10^.

An alternative argument based on ancient substructure within Africa, known as linkage disequilibrium (LD), is based not on individual alleles but on the association between stretches of the genome. Linkage disequilibrium occurs when there is a nonrandom association of alleles at two or more loci on a chromosome, such that the haplotype frequency is no longer equal to the product of the allele frequencies. In present-day human populations, the extent of LD between two single nucleotide polymorphisms (SNPs) shared with Neanderthals can be the result of either (a) “nonadmixture LD”, reflecting stretches of DNA inherited from the ancestral population of Neanderthals and modern humans as well as LD that has arisen due to bottlenecks and genetic drift in modern humans as they separated from Neanderthals, or (b) “admixture LD”, reflecting stretches of genetic material resulting from gene flow from Neanderthals into modern humans (Sankararaman et al 2012, pages 1–2).

Sankararaman et al (2012) measured the extent of LD in the genomes of present-day Europeans and found that the last gene flow from Neanderthals into Europeans likely occurred between 37–86 Kya and most likely between 47–65 Kya, which is “too recent to be consistent with the “ancient African population structure” scenario” … and “strongly supports the hypothesis that at least some of the signal of Neanderthals being more closely related to non-Africans than to Africans is due to recent gene flow” (Sankararaman et al 2012, pages 6–7). Whilst Sankararaman et al (2012) accepted Eriksson and Manica’s demonstration, using a spatially explicit model and approximate Bayesian computation, that ancient substructure can also account for the observation from D-statistics (Eriksson and Manica 2012, pages 13956–13958), they claim that both this new approach and that of Yang et al (2012) show that “ancient substructure alone cannot explain these signals”.

### Relative proportions of derived alleles in the present-day European genome

Table 3 shows the calculation of the relative proportions of derived alleles of Neanderthals versus sub-Saharan Africans versus the third archaic ancestor in the European genome. Derived Neanderthal alleles in the European genome are represented by allele pattern BABA; African derived alleles by BBAA (or CCAA); and derived alleles of the third archaic ancestor by BAAA + BACA + BCAA + BCDA (or DAAA). The corresponding allele counts are 157,935 Neanderthal derived alleles; 375,289 African derived alleles; and 630,343 third archaic ancestor derived alleles. The corresponding percentages of the European genome are 0.0112% Neanderthal; 0.0266% African; and 0.0447% third archaic ancestor, respectively. The relative proportions of derived alleles in the 0.0826% of the European genome that is not shared with the common ancestor of humans and chimpanzee are 13.6% Neanderthal, 32.3% sub-Saharan African and 54.2% third archaic ancestor.

Analysis of the allele counts from the alignment of the 45 Kya fossil from Ust’-Ishim in western Siberia, kindly provided to the author by Qiaomei Fu and Janet Kelso, shows similar relative proportions of 14.3 % Neanderthal, 31.7% sub-Saharan African and 54.0% third archaic ancestor, suggesting that this individual was closely related to the ancestor of present-day Europeans (Table 3 below; Fu et al 2014). Morphological analysis of the Ust’-Ishim femur shows closer affinities to Upper Paleolithic modern humans than to Early Anatomically Modern Humans (Early AMH), consisting of Skhul and Qafzeh samples, or to Neanderthals. (Fu et al 2014, Suppl. 3.)

Analysis of the allele counts from the alignment of the 36.2 Kya Kostenki 14 (Markina Gora) fossil from Kostenki-Borshchevo in European Russia, kindly provided to the author by Martin Sikora and Eske Willerslev, also shows similar relative proportions of 16.2% Neanderthal, 33.1% sub-Saharan African and 50.8% third archaic ancestor (Table 3 below; Seguin-Orlando et al 2014).

The similarity between the relative proportions of derived alleles in the genomes of these two archaic individuals and those of present-day Europeans, represented by the French individual, indicates a common origin in an admixed population prior to 45 Kya, with no subsequent major contribution to the European genome.

The relative proportion of Neanderthal derived alleles of 13.6% differs significantly from the proportion of Neanderthal ancestry in present-day Europeans of between 1.3% and 2.7% reported in Green et al (2010), and of between 1.48% and 1.96% reported in Prüfer et al (2014). This proportion is simply the relative proportion of derived alleles in the present-day European genome. There is no attempt in these computations to infer the history of these populations, nor does it require the use of coalescent theory or simulations. However, it is extremely difficult to explain how between 1.3% and 2.7% of Neanderthal ancestry, however defined, could result in a 13.6% relative proportion of Neanderthal derived alleles in the European genome.

The analysis in Appendix A identifies an error in the derivation of Reich and Patterson’s admixture estimators, and shows how this error resulted in a Neanderthal admixture estimate of between 1.3% and 2.7% rather than 29.6% in the two population model or 13.6% in the three population model. It also demonstrates that the admixture estimators which they actually used do not correspond to the admixture fractions in either the two population or three population admixture equations, or to their definitions of Neanderthal admixture^11^.

### A new model of human dispersal

This analysis of the genetic data indicates the existence of a third archaic ancestor of present-day Europeans, which raises the question of where this third lineage evolved. A possible candidate, indicated in the paleoanthropological and paleolithic record, is a Eurasian lineage in the Levant, which admixed with Neanderthals between 250–55 Kya as they expanded eastward, and subsequently with members or descendants of mtDNA haplogroup L3 after their emergence from Africa between 84–63 Kya. mtDNA haplogroups N, R, U, U2, U8 and JT descended from mtDNA haplogroup L3 in the Arabian Peninsula or the Levant between 63–50 Kya, possibly as a result of admixture between genetically distant humans (Soares et al 2010). It is conceivable that members of these populations, with a morphology similar to present-day *Homo sapiens*, expanded westward into Europe along the Danube and Mediterranean coast and replaced the already dwindling Neanderthal populations between 50–40 Kya, rather than sub-Saharan Africans, generally referred to as Anatomically Modern Humans (AMH).

This paper does not address the origin of other present-day populations, but the genetic, archaeological and anthropological evidence suggests radiations in all directions from a basal admixed Eurasian population in the Levant between 55–50 Kya, in addition to earlier expansions eastward from the Arabian Peninsula by members of mtDNA haplogroups L3, M, N and R along the southern route into South East Asia, Melanesia and Australia. These expansions appear to have resulted in admixture with other archaic lineages in East Asia, including Denisovans.

Ancestors of the Ust’-Ishim individual, a member of mtDNA haplogroup R, probably went northeast from the Levant into western Siberia around 47 Kya (Fu et al 2014). Ancestors of the Kostenki 14 individual, a member of mtDNA haplogroup U2, probably moved northward from the Levant into the Central European Plain between 40–36 Kya (Seguin-Orlando et al 2014).

### Conclusion

The most parsimonious explanation of the allele counts is the existence of a third archaic ancestor of Europeans, probably in the Levant, which diverged from the ancestor of Africans around 900 Kya, following an early split between the ancestors of Africans and Neanderthals around 1.4 Mya. This, together with subsequent admixture between the third archaic ancestor and Neanderthals around 250–55 Kya and with newly emerged ancestors of Africans between 63–50 Kya, provides a highly coherent explanation of all 15 allele counts in the Altai alignment that is consistent with the archaeological and anthropological record. Without this assumption, these allele counts are entirely inexplicable.

Analysis of the allele counts attributable to the three archaic ancestors of present-day Europeans shows that the relative proportions of derived alleles in the 0.0826% of the European genome that is not shared with the common ancestor of humans and chimpanzee are 13.6% Neanderthal, 32.3% sub-Saharan African and 54.2% third archaic ancestor (Table 3).

Applying the same analysis to the San genome, the relative proportions of derived alleles in the sub-Saharan African genome are 87.7% African and 12.3% Neanderthal (Table 4). This is compatible with the greater genetic isolation of sub-Saharan Africans, as might be expected from their geographical location and the obstacles to outward or inward migration.

## Appendix A

### Mathematical error in the derivation of estimators of admixture proportions

The two published estimators of admixture proportions in the European genome do not represent the admixture fraction *f* in the two population admixture equation *E* = *fN_A_* + *(1−f) A_A_,* where E is the genome of the present-day European population, N_A_ is the Neanderthal genome that contributed alleles to the present-day European, and A_A_ is the genome of the sub-Saharan African ancestral population of present-day non-Africans, because (i) the substitution of the S statistic and F_4_ statistic in the two population admixture model results in a value for *f* which is not equal to the admixture fraction; and (ii) a three population model is required if there is a third archaic ancestor of Europeans. Consequently, the resulting admixture estimators do not measure the proportion of Neanderthal derived alleles in the European genome.

In particular, the admixture estimator actually used by David Reich and Nick Patterson does not correspond to their definition of Neanderthal admixture as the proportion of lines of descent or probability that the line of descent, will pass through the Neanderthal population or lineage, or the proportion that traces its genealogy through the Neanderthal side of the phylogenetic tree. Substitution of the alleles in each genome or the aligned alleles for each ancestral population in a three population admixture equation provides a valid estimator of admixture proportions based on the relative contributions of derived alleles.

The two published estimators of the admixture proportion

A. *f* = *E[S(E, A, N_1_, C)]/ E[S(N_A_, A, N_1_, C)]* = *[Σ{i* = *1,n}[C_BABA_(i) − C_ABBA_(i)]]^E^ / [Σ{i* = *1,n}[C_BABA_(i) − C_ABBA_(i))]]^NA^*

The derivation of this admixture estimator was provided by Nick Patterson and David Reich in Green et al (2010), Suppl. 18. A genealogical derivation of the D and S statistics based on admixture graphs and coalescence theory, which was later extended to derive the *f*_4_-ratio, was provided by Eric Durand, Philip Johnson, Anna-Sapfo Malaspinas and Montgomery Slatkin in Green et al (2010), Suppl. 19; by Nick Patterson, Heng li, Swapen Mallick and David Reich in Reich et al (2010), Suppl. 8; by Eric Durand and Montgomery Slatkin in Reich et al (2010), Suppl. 11; and by Eric Durand, Nick Patterson, David Reich and Montgomery Slatkin in Durand et al (2011).

In Green et al (2010), David Reich and Nick Patterson used the difference between the allele counts for allele patterns BABA and ABBA in the European-African-Neanderthal-Chimpanzee alignment, where the European and Neanderthal share derived alleles in BABA and the African and Neanderthal share derived alleles in ABBA, to determine whether Neanderthals were more closely related to non-Africans than to Africans. They defined the D statistic as *D(E, A, N_1_, C)* = *[Σ{i* = *1,n}[C_BABA_(i) − C_ABBA_(i))]]^E^/ [Σ{{i* = *1,n}[C_BABA_(i)* + *C_ABBA_(i)]]^E^*, where E represents the present-day European population, A represents the present-day sub-Saharan African population, N_1_ represents one Neanderthal sample, and C represents the present-day Chimpanzee population, and C_BABA_ and C_ABBA_ are indicator variables, which can be 0 or 1 depending on whether an ABBA or BABA pattern is seen at base *i* (Green et al 2010, Suppl. 15). For the alignment French-San-Vindija Neanderthal-Chimpanzee, D = 0.042 or 4.2% (Green et al 2010, Suppl. 15, table S48).

The allele counts for the BABA and ABBA allele patterns were then used to estimate the admixture proportion of Neanderthal alleles in present-day Europeans, by substituting the numerator of the D statistic, the S statistic, *S(X, A, N_1_, C)* = *Σ((i* = *1,n}[C_BABA_(i) − C_ABBA_(i)],* for each genome in the two archaic ancestor admixture equation *E* = *fN_A_* + *(1−f) A_A_,* where *S(X, A, N_1_, C)* measures the relative rate of matching of samples X and A to the Neanderthal sample N_1_ which was sequenced.

This results in *E[S(E, A, N_1_, C)]* = *(f) E[S(N_A_, A, N_1_, C)]* + *(1−f) E[S(A_A_, A, N_1_, C)].*

Setting *E[S(A_A_, A, N_1_, C)]* = 0 because *A_A_* and *A* form a clade relative to N_1_ and C; *E[S(E, A, N_1_, C)]* = *(f) E[S(N_A_, A, N_1_, C)].*

Then *f= E[S(E, A, N_1_, C)]/ E[S(N_A_, A, N_1_, C)]* = *[Σ{{i* = *1,n}[C_BABA_(i) − C_ABBA_(i)]]^E^/ [Σ{{i* = *1,n}[C_BABA_(i) − C_ABBA_(i)]]^NA^,* where *[Σ{{i* = *1,n}[C_BABA_(i) − C_ABBA_(i)]]^E^* and *[Σ{{i* = *1,n}[C_BABA_(i) − C_ABBA_(i)]]^NA^* measure the relative rate of matching of samples E and A, and N_A_ and A, respectively, to the Neanderthal sample N_1_ which was sequenced.

B. *f*_4_-ratio, 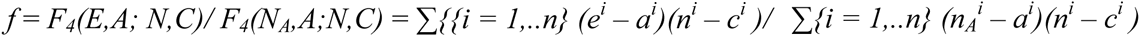

The derivation of the *f*_4_-ratio admixture estimator was provided by David Reich and Nick Patterson in Reich et al (2009), Suppl. 5; by Nick Patterson, Heng li, Swapen Mallick and David Reich in Reich et al (2010), Suppl. 8; by Eric Durand and Montgomery Slatkin in Reich et al (2010), Suppl. 11; by Nick Patterson and David Reich in Patterson et al (2012): 1072–1073 and Appendix A; by Nick Patterson, Swapan Mallick and David Reich in Meyer et al (2012), Note 11: 42; and by Nick Patterson, Swapan Mallick and David Reich in Prüfer et al (2014), Suppl. 14.

The *f*_4_-ratio was derived by substituting the F_4_ statistic, *F_4_(X,A;N,C)* = *Σ{i* = *1,…n}(x^i^ − a^i^)(n^i^ − c^i^),* for each genome in the two archaic ancestor admixture equation *E* = *fN_A_* + *(1−f)A_A_,* where E is the genome of the present-day European population, N_A_ is the Neanderthal genome that contributed alleles to the present-day European population, A_A_ is the genome of the sub-Saharan African ancestral population of present-day non-Africans, C is the genome of the present-day chimpanzee population, *F_4_(X,A;N,C)* measures the overlap between the drift paths of two admixing populations, and x*^i^*, *a^i^*, *n^i^*, *c^i^* are the corresponding allele frequencies.

This results in *F_4_(E,A; N,C)* = *(f) F4(N_A_,A;N,C)* + *(1−f) F_4_(A_A_,A;N,C).*

Setting *F_4_(A_A_,A;N,C)* = 0, because *A_A_* and *A* are in a clade relative to N and C, results in *F_4_(E,A;N,C)* = *(f) F_4_(N_A_,A;N,C).*

Then, 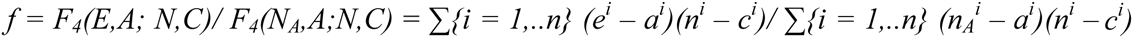 is seen to represent the overlap between drift paths between the two admixing populations on each side of the phylogeny, where *e^i^*, *a^i^*, *n^i^*, *c^i^* and 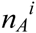 are the corresponding allele frequencies in each population (Reich et al 2009, Suppl. 5; Patterson et al 2012:1072–1073 and Appendix A).

The genealogical derivation based on admixture graphs and coalescence theory assumed that with probability *f* the E lineage was derived from a lineage N and that with probability (1-*f*) the E lineage was not derived from a lineage N but traced its lineage through the P(_1_,_2_) = P(_E_,_A_) side of the phylogeny, and that the probability of allele patterns BABA and ABBA equals the probability of the appropriate topology times the probability of coalescence, times the branch length, times the mutation rate.

#### Substitution of the S statistic and the F_4_ statistic in the admixture equation

The first question is what does the admixture equation *E* = *fN_A_* + *(1−f) A_A_* mean. What does it mean to equate one population as a linear admixture of two other populations or one genome as a linear admixture of two other genomes? It clearly does not mean that we simply add individuals from two ancestral populations, e.g. A_A_ and N_A_ in particular proportions and the result is the present-day population, E. In this case, we are defining a fraction, *f*, of the alleles in the European genome as deriving from the genome of the Neanderthal ancestor and the residual fraction (1-f) as deriving from the genome of the African ancestor. It is a vector equation between the vectors *E*, *N_A_* and *A_A_* with coefficients equal to 1 and 0 or 0 and 1 at each site on the genome. *f* is the sum of the coefficients for the vector *N_A_* normalized by the total number of sites.

In general, *f* in the admixture equation *x* = *fy* + *(1-f)z,* is only valid as an admixture proportion under certain conditions. This equation is really two equations, *x* = *ay* + *bz* and *a* + *b* = *1.* The conditions for *f* to be a valid admixture proportion in this equation include the following:

1. The values for the variables, x, y and z, must be additive, which means that they must be expressed in common units; e.g. pints, pounds, or number of alleles;
2. The values of the variables, x, y and z, must be normalized to be the same size in these units; e.g. 1 pint, 1 pound, 1 genome or a fixed number of alleles; otherwise *f* and *(1-f)* will reflect the values of the variables rather than the proportions in the dependent variable;
3. The independent variables, y and z, must both be non-zero unless *f* = 1 or 0 (no admixture) so that *a* + *b* = *1*; otherwise, for example, if z = 0, *f* in *x* = *fy* does not represent the admixture proportion unless *f* = 1, as there is no *(1-f)* to complete the mixing equation and no condition on *f*.

In derivation A, substitution of the S statistic *S(X, A, N_1_, C)* = *Σ{i* = *1,n}[C_BABA_(i) − C_ABBA_(i)],* for each population X in the two archaic ancestor admixture equation *E* = *fN_A_* + *(1−f) A_A_,* and elimination of the term *(1-f) A_A_* because *S(A_A_, A, N_1_, C) =* 0, clearly does not satisfy the conditions for *f* to be a valid admixture proportion. For *f* to be a valid admixture proportion in this equation, the S statistic *S(X, A, N_1_, C)* = *Σ{i* = *1,n}[C_BABA_(i) − C_ABBA_(i)]* for each population would need to be valid variables in the admixture equation. However, it is clear that they are not: (i) is satisfied as numbers of bases *i* (sites or alleles) are additive; (ii) is not satisfied; the value of the S statistic is not the same for each population; and (iii) is not satisfied unless *f* = 1 (no admixture) because *E[S(A_A_, A, N_1_, C)]* = 0.

Consequently, rather than being derived from *E* = *fN_A_* + *(1−f) A_A_,* we are left with the admixture proportion being defined as *f^*^* = *E[S(E, A, N_1_, C)]/E[S(N_A_, A, N_1_, C)]* = *[Σ{i* = *1,n}[C_BABA_(i) − C_ABBA_(i)]]^E^/ [Σ{i* = *1,n}[C_BABA_(i) − C_ABBA_(i))]]^NA^,* where *[Σ{i* = *1,n}[C_BABA_(i) − C_ABBA_(i)]]^E^* and *[Σ{i* = *1,n}[C_BABA_(i) − C_ABBA_(i)]]^NA^* measure the relative rate of matching of samples E and A, and N_A_ and A, respectively, to the Neanderthal N_1_ which was sequenced.

Nonetheless, it is worth understanding what is being measured by this ratio. The meaning of *[Σ{i* = *1,n}[C_BABA_(i) − C_ABBA_(i)]]^E^* as the relative rate of matching of samples E and A, to N_1_ is straightforward; but it is less clear what the denominator *[Σ{i* = *1,n}[C_BABA_(i) − C_ABBA_(i)]]^N^* means when we are matching alleles of one Neanderthal against another at sites which are restricted to Neanderthal derived alleles, or how the ratio of these two terms relates to the admixture proportion *f* in the admixture equation *E* = *fN_A_* + *(1−f) A_A_*.

The D statistic is defined as D = *[Σ{i* = *1,n}[C_BABA_(i) − C_ABBA_(i)]]^E^/ [Σ{i* = *1,n}[C_BABA_(i)* + *C_ABBA_(i)]]^E^.*

Substituting *[Σ{i = 1,n}[C_BABA_(i) – C_ABBA_(i)]]^E^ = Dx [Σ{i = 1,n}[C_BABA_(i) + C_ABBA_(i)]]^E^* in *f = [Σ{i = 1,n}[C_BABA_(i) − C_ABBA_(i)]]^E^/ [Σ{i = 1,n}[C_BABA_(i) – C_ABBA_(i)]]^NA^,* gives *f = D x [Σ{i = 1,n}[C_BABA_(i) + C_ABBA_(i)]]^E^ / [Σ{i = 1,n}[C_BABA_(i) − C_ABBA_(i)]]^NA^*.

Substituting D = 0.042 from Table S48 in Green et al (2010), *f* = 0.042 *x [Σ{i* = *1,n}[C_BABA_(i)* + *C_ABBA_(i)]]^E^ / [Σ{i* = *1,n}[C_BABA_(i)* − *C_ABBA_(i)]]^NA^*. (Green et al 2010, Suppl. 15, table S48).

Substituting the observed allele counts in the numerator, *[Σ{i* = *1,n}[C_BABA_(i)* + *C_ABBA_(i)]]^E^* = BABA + ABBA = 157,935 + 143,385 = 301,320.

Ignoring *C_ABBA_(i)* in the denominator *[Σ{i* = *1,n}[C_BABA_(i)* − *C_ABBA_(i)]]^NA^* because this is likely to be small compared with *C_BABA_(i)* for N_A_AN_1_C, the denominator reduces to *[Σ{i* = *1,n}[C_BABA_(i)* − *C_ABBA_(i)]]^NA^* = *[Σ{i* = *1,n}[C_BABA_(i)]^NA^*.

Substituting the observed allele counts in the denominator, recognizing that if the two Neanderthal samples are broadly similar, the number of alleles in NA matching those in N1 is approximately equal to the total number of Neanderthal derived alleles in the Neanderthal genome, *[Σ{i* = *1,n}[C_BABA_(i)* − *C_ABBA_(i)]]^NA^* = *[Σ{i* = *1,n}[C_BABA_(i)]^NA^* = AABA + BABA + ABBA = 755,369 + 157,935 + 143,385 = 1,056,689.

Consequently, the admixture estimator defined as *f^*^* = *E[S(E, A, N_1_, C)]/ E[S(N_A_, A, N_1_, C)]* = *[∑{i* = *1,n}[C_BABA_(i)* − *C_ABBA_(i)]]^E^/ [∑{i* = *1,n}[C_BABA_(i)* − *C_ABBA_(i)]]^NA^* = 0.042 × 301,320/1,056,689 = 1.2%. This value is a lower bound because with less than 100% matching of Neanderthal derived alleles between the two Neanderthal samples N_A_ and N_1_, or if *C_ABBA_(i)* is non-zero, the value of the denominator will be reduced and this percentage will be increased. This is then not far from the estimates of between 1.3% and 2.7 % reported in Green et al (2010), Suppl. 18: 161; 3.0% in Reich et al (2010), Suppl. 8: 55; and between 1.48% and 1.96% in Prüfer et al (2014), Suppl. 14: 128. This also demonstrates the relationship between this estimator *f^*^* and the D statistic.

*f^*^* = *E[S(E, A, N_1_, C)]/ E[S(N_A_, A, N_1_, C)]* is clearly not the admixture proportion *f* in the admixture equation *E* = *fN_A_* + *(1*−*f) A_A_.* The ratio was described in Green et al (2010) as “the extent to which the European population is towards being entirely of Neanderthal ancestry”. (Green et al 2010, Suppl. 18: 159). That is not what *f* represents in the two archaic ancestor admixture equation *E* = *fN_A_* + *(1*−*f) A_A_*; nor is it the Neanderthal admixture proportion *f^N^* in a three archaic ancestor admixture equation *E* = *f^N^N_A_* + *f^A^A_A_* + *(1*−*f^N^*−*f^A^)O_A_*, where O_A_ is the genome of the third archaic ancestor of Europeans. Nor does the admixture estimator actual used by David Reich and Nick Patterson correspond to their definition of Neanderthal admixture as the proportion of lines of descent or probability that the line of descent, will pass through the Neanderthal population or lineage, or the proportion that traces its genealogy through the Neanderthal side of the phylogenetic tree.

If the admixture equation is restricted to derived alleles of the Neanderthal and sub-Saharan African ancestors in the European genome, in the two archaic ancestor model, the proportion *f* of Neanderthal derived alleles relative to the total number of derived alleles of the Neanderthal and African ancestors in the European genome is *f* = BABA/(BABA + BBAA) = 29.6% (Table 3). In contrast, *f^*^* is approximately equal to the proportion of the derived alleles with allele patterns BABA and ABBA relative to the total number of Neanderthal derived alleles in the Neanderthal genome = <u>0.042</u> × (BABA + ABBA)/ (<u>AABA</u> + BABA + ABBA) = 1.2%. The numerical difference between these two estimates is due primarily to the multiplication by the D statistic in the computation of *f^*^*, and secondarily to the addition of ABBA to the numerator.

In the three archaic ancestor model, the proportion of Neanderthal derived alleles in the European genome relative to the total number of derived alleles of the three archaic ancestors in the European genome is *f* = BABA/(BAAA + BABA + BBAA) = 13.6% (Tables 2 and 3).

In derivation B, substitution of the F_4_ statistic, *F_4_(X,A;N,C)* = *∑{i* = *1,…n} (x^i^* − *a^i^)(n^i^* − *c^i^),* for each population X in *E* = *fN_A_* + *(1*−*f) A_A_,* and elimination of the term *(1-f) A_A_* because *F_4_(A_A_,A;N,C)* = 0, clearly does not satisfy the conditions for *f* to be a valid admixture proportion. For *f* to be a valid admixture proportion in this equation, the F_4_ statistic *F_4_(X,A;N,C)* = *∑{i* = *1,…n} (x^i^* − *a^i^)(n^i^* − *c^i^)* for each population would need to be valid variables in the admixture equation. However, it is clear that they are not: (i) is satisfied as the allele frequencies are additive; (ii) is not satisfied; the value of the F_4_ statistic is not the same for each population; and (iii) is not satisfied unless *f* = 1 (no admixture) because *F_4_(A_A_,A;N,C)* = 0.

Consequently, rather than being derived from *E* = *fN_A_* + *(1*−*f) A_A_*, we are left with the admixture proportion being defined as 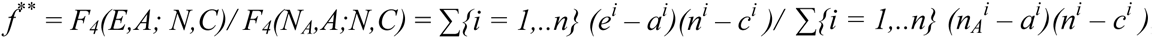, where *e^i^*, *a^i^*, *n^i^*, *c^i^* and 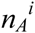 are the corresponding numbers of alleles in the drift paths between A and N and N_A_ and N on each side of the phylogeny. As with *f^*^* = *E[S(E, A, N_1_, C)]/ E[S(N_A_, A, N_1_, C)]*, the *f*_4_-ratio for a Eurasian was interpreted “intuitively … as measuring how far of the way a Eurasian population is toward having the allele frequency patterns with Africans, Denisovans and chimpanzee that is characteristic of a 100% Neanderthal” (Reich et al 2010, Suppl. 8: 55). As before, that is not what *f* represents in the population model *E* = *fN_A_* + *(1*−*f) A_A_*, nor is it the Neanderthal admixture proportion *f^N^* in a three archaic ancestor population model *E* = *f^N^N_A_* + *f^A^A_A_* + *(1*−*f^N^*−*f^A^)O_A_*.

The alleles in a genome or the aligned alleles constitute a valid variable in the admixture equation

Although the substitutions of S and F4 statistics in the two population admixture equation *E* = *fN_A_* + *(1*−*f) A_A_* have been demonstrated as invalid, this equation appears to be a reasonable description for the admixture of two archaic populations; and *E* = *f^N^N_A_* + *f^A^A_A_* + *(1*−*f^N^*−*f^A^)O_A_* appears to be a reasonable description for the admixture of three archaic populations. As we have seen above, for *f* to be a valid estimator of the admixture proportion, the population variable substituted for the three populations must be of constant size. It could be the entire genome of each population, or it could be the alleles aligned in each population. Condition (i) is satisfied because numbers of alleles are additive; (ii) is satisfied because the same number of alleles for the same sites have been selected for each population in the alignment; and (iii) is satisfied because there is a set of alleles for each population. If the variables in the admixture equation are the aligned alleles for each population, *f* or *f^N^* will be the relative proportion of Neanderthal alleles in the European genome.

### Conclusion

The two published estimators of admixture proportions in the European genome *f* = *E[S(E, A, N_1_, C)]/ E[S(N_A_, A, N_1_, C)]* and the *f*_4_-ratio, *f* = *F_4_(E,A; N,C)/ F_4_(N_A_,A;N,C)* do not represent the admixture fraction *f* in the two population admixture equation *E* = *fN_A_* + *(1*−*f) A_A_* because (i) the substitution of the S statistic and F_4_ statistic in the two population admixture equation results in a value for *f* which is not equal to the admixture fraction; and (ii) a three population admixture equation is required if there is a third archaic ancestor of Europeans. Consequently, the resulting admixture estimators do not measure the proportion of Neanderthal derived alleles in the European genome.

Moreover, it is extremely difficult to explain how between 1.3% and 2.7% of Neanderthal ancestry, however defined, could result in a 13.6% relative proportion of Neanderthal derived alleles in the European genome. Consequently, the estimates of Neanderthal ancestry in the European genome of between 1.3% and 2.7% in Green et al (2010), Reich et al (2010) and Prüfer et al (2014) must be rejected. The alleles in a genome or the aligned alleles constitute a valid variable in the admixture equation.

## Acknowledgements

The author would like to thank Nick Patterson and David Reich for assistance in providing the unpublished allele counts for the French-San-Altai Neanderthal-Chimpanzee alignment; Qiaomei Fu, Janet Kelso and David Reich for assistance in providing the Ust’Ishim-San-Altai Neanderthal-Chimpanzee allele counts; and Martin Sikora, Eske Willerslev and Robert Foley for assistance in providing the Kostenki-San-Altai Neanderthal-Chimpanzee allele counts. The author would also like to thank David Reich, Nick Patterson, Montgomery Slatkin, Michael Walker, Paul O’Higgins, Leslie Aiello and three anonymous reviewers for their helpful comments on earlier versions of this paper.

## Footnotes

1. This paper does not address Denisovan admixture as there is no evidence of Denisova or their DNA in Europe, and the available allele counts for Denisova do not permit the same type of analysis because Denisovan DNA occurs alongside Neanderthal DNA in Asian samples and is difficult to disentangle.

2. The human mutation rate has recently been reduced from 1.0 × 10^−9^ bp per year to around 0.5 × 10^−9^ bp per year. The 1000 Genomes Project Consortium in 2010 calculated mutation rates based on de novo germline mutations for the present-day human samples of 1.2 × 10^−8^ bp per generation for Europeans and 1.0 × 10^−8^ bp per generation for Africans compared with 0.5 × 10^−9^ bp per year for the archaic ancestors assumed in this paper. These correspond to generations of 24 and 20 years respectively (1000 Genomes Project Consortium 2010, page 1,068 and Suppl. Info. 12.4).

3. Patterson and Reich confirmed that the error rate in these allele counts was relatively small, stating that “assuming that the mutation rates have been constant on both French and San lineages, the difference in the rates of ABAA and BAAA counts (647,939 and 629,542 respectively) must be due to a difference in the error rate of the sequencing for French and San” (Green et al 2010, SOM 15, page 137). This difference was used to estimate the error rate, and to validate their analysis based on the difference between the ABBA and BABA allele counts. In the testing of filters in Suppl. Info. 6a in Prüfer et al, the mutations on each lineage leading to Africans, Europeans, Neanderthal and Denisova were compared with the mutations on the human reference genome and the ratios were shown to be close to 1 after allowing for branch shortening due to difference in ages of death (Prüfer et al 2014, pages 45–6, Suppl. Info. 6a). This assumes equal mutation rates on all lineages.

4. Levantine Mousterian artifacts dated between 250–40 Kya, at many sites in association with Neanderthal or modern human fossils with Neanderthal affinities, have been found at Dederiyeh Cave, Afrin Valley, northwestern Syria; at Hummal, Umm el Tlel and Nadaouiyeh ain Askar in the El Kowm basin, in central Syria; at Jerf al-Ajla cave, in the Palmyra basin, central Syria; at Yabrud I Rockshelter and Yabrud II Rockshelter, in the Skifta Valley, Syria; at Ras el Kelb, north of Beirut, Lebanon; at Bezez Cave, Aadloun, in southern Lebanon; at Nahal Mahanayeem Outlet, on the eastern bank of the Upper Jordan River, northern Israel; at Amud Cave, 5 km northwest of the Sea of Galilee, northern Israel; at Zuttiyeh Cave, near the Sea of Galilee, Israel; at Hayonim Cave, Upper Galilee, Israel; at Jebel Qafzeh Rockshelter, Precipe Mountain, Lower Galilee, Israel; at Es Skhul Cave, Nahal Me’arot canyon near Haifa, Israel; at Tabun Cave, Jamal Cave and Misliya cave, Mount Carmel, Israel; at Rosh Ein Mor, Central Negev, Israel; at ‘Ain Dfla Rockshelter, Wadi Ali, west-central Jordan; and at Jebel Qattar, in the Jubbah Palaeolake, Nefud Desert, northern Saudi Arabia (Nishiaki et al 2012, Eurasian Prehistory; Institute for Prehistory and Archaeological Science 2006, University of Basel; Schwarcz et al 2001, Paleorient; Solecki et al 1986, Paleorient; Pastoors et al 2008, Paleorient; Copeland 1982, Paleorient; Kalbe et al 2014, Quaternary International; Hovers et al 1995, Paleorient; Bar-Yosef et al 1974, Paleorient; Mercier et al 2007, Journal of Archeological Science; Bar-Yosef Mayer et al 2009, Journal of Human Evolution; Grun et al 2005, Journal of Human Evolution; Jelinek 1982, Science; Tsatskin et al 1994, Paleorient; Weinstein-Evron et al 2012, PaleoAntropology; Rink et al 2003, Journal of Archaeological Science; Mustafa et al 2007, Eurasian Prehistory; Petraglia et al 2011, Quaternary Science Reviews).

5. Homo rhodesiensis cannot be the ancestor of Homo neanderthalensis as the DNA indicates a much earlier separation between the ancestors of Africans and the ancestors of Neanderthals. Interestingly, early admixture between the African lineage and the Neanderthal lineage (in the form of Homo heidelbergensis), resulting in an “anatomically modern human” (AMH) morphology in Africa, might be a precursor to the later admixture of the African lineage with a hybrid of the third archaic ancestor lineage and Neanderthals to create another anatomically modern human, present-day humans or Homo sapiens, in Eurasia. Even though it is claimed in Prüfer et al (2014), that present-day sub-Saharan Africans share more derived alleles with Neanderthals than with Denisovans, coalescent simulations based on a model in which Neanderthals admixed with the ancestors of sub-Saharan Africans after their split from Denisova around 600 Kya, did not predict an increase in this signal at sites with a higher frequency of African derived alleles. (Prüfer et al 2014, Suppl. 16a.) However, this would not be expected if the admixture occurred around 600 Kya with a group of early Homo heidelbergensis, prior to the split of Denisovan ancestors from another group of Neanderthal ancestors, for example, which had headed east into the Altai Mountains at around the same time. In any event, the ABBA allele counts in both Green et al (2010) and Prüfer et al (2014) indicate that admixture occurred between the ancestors of both the Vindija and Altai Neanderthals and ancestors of sub-Saharan Africans (Table 1).

6. Indications of the existence of an unknown other archaic ancestor of Denisovans were also reported in Prüfer et al (2014), which found that sub-Saharan Africans share more derived alleles with Neanderthals than with Denisovans and that this signal grows stronger for alleles that occur at 100% frequency (i.e. are fixed) in Africans (Prüfer et al 2010, pages 46–47 and Suppl. 16a, page 139). This was interpreted as being best explained by gene flow from an unknown archaic population into Denisovans, which diverged from ancestors of Africans and Neanderthals around 0.9–1.4 Mya. The estimates above for divergence dates fall within this range, so this could have been the third archaic ancestor.

7. There is archaeological and anthropological evidence of Oldowan I artifacts and early human fossils dated from 1.4–1.0 Mya at Ubeidiya, central Jordan Rift Valley, 3 Km south of Sea of Galilee, Israel (Shea et al 1999, Journal of Israel Prehistoric Society). There is also archaeological and anthropological evidence of Oldowan I artifacts and early human fossils dated from 1.4–1.2 Mya at Barranco Leon and Fuente Nueva 3 in the Orce region, Guadix Baza Basin, Granada, Spain and at Sima del Elefante, Trinchera del Ferrocarril, Atapuerca, Burgos, Castille, Spain (Toro-Moyano et al 2013, Journal of Human Evolution; Ribot et al 2015, Current Anthropology; Barsky et al 2015, Quaternary International; Rodriguez et al 2011, Quaternary Science Reviews).

8. There is archaeological and anthropological evidence of Acheulian artifacts dated from 1.0 Mya-780 Kya at Bizat Ruhama, southern coastal plain, Northern Negev, Israel; at Hummal, El Kowm Oasis, northeast of Palmyra, central Syria; and at Evron Quarry, northern coastal plain of Israel (Belmaker et al 2002, Journal of Human Evolution; Zaidner et al 2010, PaleoAnthropology; Zaidner 2013, PLOS ONE; Le Tensorer et al 2008, Basle Symposium; Richter et al 2008, Basle Symposium; Ron et al 2003, Journal of Human Evolution; Tchernovet al 1994, Quaternary Research). At Hummal, Umm el Tlel andNadaouiyeh ain Askar in the El Kowm basin, central Syria; at Yabrud I in the Skifta Valley, Syria; `at` Bezez Cave, Aadloun, southern Lebanon; and at Tabun, Mount Carmel, Israel, the Early and Middle Acheulian was followed by long sequences including the Acheulo-Tayacian, Tayacian, Late Acheulian, Acheulo-Yabrudian, Yabrudian, and Hummalian, before the arrival of the Levalloiso-Mousterian around 250 Kya (Institute for Prehistory and Archaeological Science 2006, University of Basel; Solecki et al 1986, Paleorient; Copeland 1982, Paleorient; Jelinek 1982, Science). The site of Gesher Benot Ya’aqov on the shore of paleolake Hula, and the associated sites with large flake Acheulian artifacts at Qana Oasis, near Jubbah, in the Nefud Desert, northern Arabia; at Saffaqah, in the Nedj peneplain, central Saudi Arabia; and at Wadi Fatimah, near the Red Sea, Saudi Arabia, are assumed to represent a separate exodus from Africa around 800 Kya and a dead-end in the Levant (Sharon 2010, Quaternary International; Shipton et al 2014, PaleoAnthropology; Petraglia et al 2009, The Evolution of Human Populations in Arabia).

9. Analysis of polymorphic SNP (Single Nucleotide Polymorphism) sites in the present-day human genome based on The International HapMap Consortium data indicated that derived allele frequencies for Europeans were between 0–5% for about 30% of the polymorphic SNPs, 5–10% for about 15% of the SNPs, and between 10–100% for the remaining 55% of the SNPs; each 10% allele frequency interval covering about 1.5–4.75% of the derived alleles, and averaging a derived allele frequency of around 28.5% for all polymorphic SNPs. The derived allele frequencies for Africans were between 0–5% for 22% of the SNPs, 5–10% for about 22% of the SNPs, and between 10–100% for the remaining 56% of the SNPs; each 10% allele frequency interval covering about 2–4% of the derived alleles, and averaging a derived allele frequency of around 25.9% for all polymorphic SNPs (The International HapMap Consortium 2007, Suppl. Info. Figure 4(c)). Applying these average derived allele frequencies to The 1000 Genomes Project Consortium low coverage mean variant SNP sites per individual of 2,918,623 for Europeans and 3,335,795 for Africans (Yoruba), which included all sites with an allele frequency of 1% or higher, including rare SNPs, and 99% of the sites genotyped in HapMap II, this represents around 2,918,623 × 0.285 = 832,808 derived alleles at polymorphic sites in present-day Europeans and 3,335,795 × 0.259 = 863,971 derived alleles at polymorphic sites in present-day Africans (The 1000 Genomes Project Consortium 2010, Table 1 and pages 1062–3). Prorating these numbers of alleles for the difference between the Altai Neanderthal alignment coverage of 1.4 billion of the approximately 3 billion alleles in the human genome compared with 2.43 billion sequenced alleles in the 1000 Genomes Project Consortium analysis, the expected number of derived alleles contributed by polymorphic sites in the French individual, represented by DAAA allele counts, is around 832,808 × 1.4/2.43 = 479,807; and the expected number of derived alleles contributed by polymorphic sites in the African individual, represented by ACAA allele counts, is around 863,971 × 1.4/2.43 = 497,761. These correspond to 685 Ky and 711 Ky of mutations respectively, compared with the branch lengths for the DAAA allele count of 900 Ky and for the ACAA allele count of 864 Ky. This suggests that most mutations appear to remain polymorphic for a very long time, even if at increasingly reduced derived allele frequencies, only reaching fixation after several hundred thousand years.

10. Lohse and Frantz (2014) noted that the D-statistic is a drastic summary of genetic variation based on mutation patterns ABBA and BABA that are incongruent with the species tree, and the fact that an excess of either sites cannot be explained by incomplete lineage sorting so must reflect Neanderthal admixture (Lohse and Frantz 2014, page 1242). They compared two alternative models of divergence, recent instantaneous unidirectional admixture (IUA) and persistent structure in the ancestral population (AS), using a more powerful maximum likelihood test, which uses all polymorphic sites not just shared derived sites, to demonstrate that IUA provides a better explanation of the allele patterns. Use of this methodology to test a third, more straightforward, model based on three lineages, and two admixture events as proposed in this paper would probably provide an even better fit with the observed allele counts. Ideally, this should use the Altai Neanderthal DNA rather than the Vindija Neanderthal DNA, which Lohse and Frantz (2014) used in their test, to avoid or minimize the correction for Neanderthal singletons (Lohse and Frantz 2014, page 1246).

11. The analysis in Appendix A identifies an error in deriving the admixture estimators f = E[S(E, A, N_1_, C)]/E[S(N_A_, A, N_1_, C)] and the f_4_-ratio, f = F_4_(E,A; N,C)/F_4_(N_A_,A;N,C) used in Green et al (2010), Prüfer et al (2014) and a number of other papers. (Reich et al 2009, Suppl. S5: 43–44 and Appendix 2: 4–9; Reich et al 2010, Suppl. 8: 43–58; Suppl. 11: 49–58; Suppl. 19: 158–161; Durand et al 2011: 2248; Reich et al 2011: 517–523 and Appendix A: 523–525; Moorjani et al 2011: 2, 9, fig. 2; Meyer et al 2012, Note 11: 42–46; Patterson et al 2012: 1072–1073 and Appendix A: 1089; Wall et al 2013: 202–203; Lazaridis et al 2014, Suppl. 6: 37–38; Fu et al 2014, Suppl. 16: 92–95; Seguin-Orlando et al 2014, Suppl. S9: 21–24.) The substitution of S statistics and F_4_ statistics in the admixture equation E = fN_A_ + (1−f) A_A_ results in a value for f which is not equal to the admixture fraction. The derivation of these admixture estimators is also in error if there is a third archaic ancestor of Europeans, for which a three population model is required. The analysis in Appendix A also demonstrates that the alleles in a genome or total number of derived alleles, used in this paper, constitute a valid variable in the admixture equation.

## References

Bar-Yosef, O., Itzik Gisis. 1974. New excavation in Zuttiyeh Cave, Wadi Amud, Israel. Paleorient 2, 1: 175–180.

Bar-Yosef Mayer, Daniella E., Bernard Vandermeersch, Ofer Bar-Yosef. 2009. Shells and ochre in Middle Paleolithic Qafzeh Cave, Israel. Journal of Human Evolution 56: 307–314.

Barsky, Deborah, Robert Sala, Leticia Menendez, and Isidro Toro-Moyano. 2015. Use and re-use: Re-knapped flakes from the Mode 1 site of Fuente Nueva 3 (Orce, Andalucia, Spain). Quaternary International 361: 21–33.

Belmaker, Miriam and Eitan Tchernov. 2002. New evidence for hominid presence in the Lower Pleistocene of the Southern Levant. Journal of Human Evolution 43: 43–56.

Copeland, Lorraine. 1978. The Middle Paleolithic of Adlun and Ras el Kelb (Lebanon): First results from a study of the flint industries. Paleorient 4: 33–57.

Durand, Eric Y., Nick Patterson, David Reich, and Montgomery Slatkin. 2011. Testing for ancient admixture between closely related populations. Molecular Biology and Evolution 28(8): 2239–2252.

Eriksson, Anders, and Andrea Manica. 2012. Effect of ancient population structure on the degree of polymorphism shared between modern human populations and ancient hominins. PNAS, 109, 35, 13956–13960.

Fu, Qiaomei, Heng Li, Priya Moorjani, Flora Jay, Sergey M. Slepchenko, Aleksei A. Bondarev, Philip L. F. Johnson, et al. 2014. Genome sequence of a 45,000-year-old modern human from western Siberia. Nature 514: 445–450.

Green, Richard E., Johannes Krause, Adrian W. Briggs, Tomislav Maricic, Udo Stenzel, Martin Kircher, Nick Patterson, et al. 2010. A draft sequence of the Neanderthal genome. Science 328: 710–722.

Grun, Rainer, Chris Stringer, Frank McDermott, Roger Nathan, Naomi Porat, Steve Robertson, Lois Taylor, et al. 2005. U-series and ESR analyses of bone and teeth relating to the human burials from Skhul. Journal of Human Evolution 49: 316–334.

Higham, Tom, Katerina Douka, Rachel Wood, Christopher Bronk Ramsey, Fiona Brock, Laura Basell, Marta Camps, et al. 2014. The timing and spatiotemporal patterning of Neanderthal disappearance. Nature 512: 306–309.

Hovers, E., Y. Rak, R. Levi, and W.H. Kimbel. 1995. Hominid remains from Amud Cave in the context of the Levantine Middle Paleolithic. Paleorient 21, 2: 47–61.

Institute for Prehistory and Archaeological Science. 2006. Research on the Paleolithic of the El Kowm area (Syria). Abstract. University of Basel.

Jelinek, Arthur J. 1982. The Tabun Cave and Paleolithic Man in the Levant. Science 216: 1369–1375.

Kalbe, Johannes, Gonen Sharon, Naomi Porat, Chengjun Zhang, and Steffen Mischke. 2014. Quaternary International 331: 139–148.

Lazaridis, Iosif, Nick Patterson, Alissa Mittnik, Gabriel Renaud, Swappan Mallick, Karola Kirsanow, Peter H. Sudmant, et al. 2014. Ancient human genomes suggest three ancestral populations for present-day Europeans. Nature 513: 409–416.

Le Tensorer, Jean-Marie, Vera Von Falkenstein, Helene Le Tensorer, and Sultan Muhesen. 2008. Hummal: A very long Paleolithic sequence in the steppe of central Syria—considerations on Lower Paleolithic and the beginning of the Middle Paleolithic. Basle Symposium: 235–248.

Lohse, Konrad, and Laurent A.F. Frantz. 2014. Neandertal Admixture in Eurasia Confirmed by Maximum-Likelihood Analysis of Three Genomes. Genetics 196, 1241–1251.

Mercier, N., H. Valladas, L. Froget, J.-L. Joron, J.-L. Reyss, S. Weiner, P. Goldberg, et al. Hayonim Cave: a TL-based chronology for this Levantine Mousterian sequence. Journal of Archeological Science 34: 1064–1077.

Meyer, Matthias, Martin Kircher, Marie-Theres Gansauge, Heng Li, Fernando Racimo, Swapan Mallick, Joshua G. Schraiber, et al. 2012. A high coverage genome sequence from an archaic Denisovan individual. Science 338: 222–226.

Mikkelsen, Tarjei S., LaDeana W. Hillier, Evan E. Eichler, Michael C. Zody, David B. Jaffe, Shiaw-Pyng Yang, Wolfgang Enard, et al. 2005. Initial sequence of the chimpanzee genome and comparison with the human genome. Nature 437: 69–87.

Moorjani, Priya, Nick Patterson, Joel N. Hisrchhorn, Alon Keinan, Li Hao, Gil Atzmon, Edward Burns, et al. 2011. The history of African gene flow into Southern Europeans, Levantines, and Jews. PLOS Genetics 7(4): e1001373.

Mustafa, Mentor, Geoffrey A. Clark. 2007. Quantifying diachronic variability: The ‘Ain Difla rockshelter (Jordan) and the evolution of Levantine Mousterian technology. Eurasian Prehistory 5(1): 47–83.

Nishiaki, Yoshihiro, Yosef Kanjo, Sultan Muhesen, and Takeru Akazawa. 2012. The temporal variability of Late Levantine Mousterian lithic assemblages from Dederiyeh Cave, Syria. 2012. Eurasian Prehistory 9, 1–2: 3–27.

Pääbo, Svante. 2014. Neanderthal Man. In search of lost genomes. New York: Basic Books.

Pastoors, A, G.-C. Weniger, and J.F. Kegler. 2008. The Middle-Upper Paleolithic transition at Yabroud II (Syria). A re-evaluation of the lithic material from the Rust excavation. Paleorient 34, 2: 47–65.

Patterson, Nick, Priya Moorjani, Yontao Luo, Swapan Mallick, Nadin Rohland, Yiping Zhan, Teri Genschoreck, et al. 2012. Ancient admixture in human history. Genetics 192: 1065–1093.

Petraglia, Michael D., Abdulla M. Alsharekh, Remy Crassard, Nick A. Drake, Huw Groucutt, Adrian G. Parker, and Richard G. Roberts. 2011. Middle Paleolithic occupation on a Marine Isotope Stage 5 lakeshore in the Nefud Desert, Saudi Arabia. Quaternary Science Reviews 30: 1555–1559.

Petraglia, Michael D., Nick Drake, and Abdullah Alsharekh. 2009. Chapter 8. Acheulean Landscapes and Large Cutting Tools Assemblages in the Arabian peninsula. The Evolution of Human Populations in Arabia. Springer Science & Business Media.

Prüfer, Kay, Fernando Racimo, Nick Patterson, Flora Jay, Sriram Sankararaman, Susanna Sawyer, Anja Heinze, et al. 2014. The complete genome sequence of a Neanderthal from the Altai Mountains. Nature 505: 43–49.

Reich, David, Kumarasamy Thangaraj, Nick Patterson, Alkes L. Price, and Lalji Singh. 2009. Reconstructing Indian population history. Nature 461: 489–495.

Reich, David, Richard E. Green, Martin Kircher, Johannes Krause, Nick Patterson, Eric Y. Durand, Bence Viola, et al. 2010. Genetic history of an archaic hominin group from Denisova Cave in Siberia. Nature 468: 1053–1060.

Reich, David, Nick Patterson, Martin Kircher, Frederick Delfin, Madhusudan R. Nandineni, Irina Pugach, Albert Nin-Shan Ko, et al. 2011. Denesova admixture and the first modern human dispersals into Southeast Asia and Oceania. American Journal of Human Genetics 89: 516–528.

Ribot, Francesc, Luis Gilbert, Carles Ferrandez-Canadell, Enrique Garcia Olivares, Florentina Sanchez, and Maria Leria. 2015. Two Deciduous Human Molars from the Early Pleistocene Deposits of Barranco Leon (Orce, Spain). Current Anthropology 56, 1: 134–142.

Richter, Daniel, Thomas C. Hauck, Dorota Wojtczak, Jean-Marie Le Tensorer, and Sultan Muhesen. 2008. Chronometric age estimates for the site of Hummal (El Kowm, Syria). Basle Symposium: 249–261.

Rightmire, G. Philip. 1996. The human cranium from Bodo, Ethiopia: evidence for speciation in the Middle Pleistocene? Journal of Human Evolution 31: 21–39.

Rink, W.J., D. Richter, and H.P. Schwarcz. 2003. Age of the Middle Palaeolithic Site of Rosh Ein Mor, Central Negev, Israel: Implications for the Age Range of the Earloy Levantine Mousterian of the Levantine Corridor. Journal of Archaeological Science 30: 195–204.

Rodriguez, J, F. Burjachs, G. Cuenca-Bescos, N. Garcia, J. Van der Made, A. Perez Gonzalez, H.-A. Blain, et al. 2011. One million years of cultural evolution in a stable environment at Atapuerca (Burgos, Spain). Quaternary Science Reviews 30: 1396–1412.

Ron, Hagai, Naomi Porat, Avraham Ronen, Eitan Tchernov, and Liora K. Horwitz. 2003. Magnetostratiography of the Evron Member—implications for the age of the Middle Acheulian site of Evron Quarry. Journal of Human Evolution 44: 633–639.

Sankararaman, Sriram, Nick Patterson, Heng Li, Svante Paabo, and David Reich. 2012. The Date of Interbreeding between Neandertals and Modern Humans. PLOS Genetics 8, 10: 1–9.

Seguin-Orlando, Andaine, Thorfinn S. Korneliussen, Martin Sikora, Anna-Sapfo Malaspinas, Andrea Manica, Ida Moltke, Anders Albrechtsen, et al. 2014. Genomic structure in Europeans dating back at least 36,200 years. Science 346: 1113–1118.

Schwarcz, H.P., P.j. Julig, W.j. Rink, H.B. Schroeder, and D. Richter. 2001. The Middle to Upper Paleolithic Transition in the Levant and new Thermoluminescence Dates for a Late Mousterian Assemplage from Jerf-al Ajla Cave (Syria). Paleorient 27, 2; 29–46.

Sharon, Gonen. 2010. Large flake Acheulian. Quaternary International 223–224: 226–233.

Shea, John J., and Ofer Bar-Yosef. 1999. Lithic Assemblages from New (1988-1994) Excavations at ‘Ubeidiya: A preliminary Report. J. of Israel Prehistoric Soc. 28: 5–20.

Shipton, Ceri, Ash Parton, Paul Breeze, Richard Jennings, Huw S. Groucutt, Tom S. White, Nicholas Drake, et al. 2014. Large Flake Acheulean in the Nefud Desert of Northern Arabia. PaleoAnthropology: 446–462.

Soares, Pedro, Alessandro Achilli, Ornella Semino, William Davies, Vincent Macaulay, Hans-Jurgen Bandelt, Antonio Torroni, et al. 2010. The Archaeogenetics of Europe. Current Biology 20: R174–R183.

Solecki, R.S., and R.L. Solecki. 1986. A Reappraisal of Rust’s Cultural Stratigraphy of Yabroud Shelter I. Paleorient 12,1: 53–59.

Tchernov, Eitan, and Liora Kolska Horwitz. 1994. The Faunal Remains from Evron Quarry in Relation to Other Lower Paleolithic Hominid Sites in the Southern Levant. Quaternary Research 42: 328–339.

The 1000 Genomes Project Consortium. 2010. A map of human genome variation from population-scale sequencing. Nature 467: 1061–1073.

The International HapMap Consortium. 2007. A second generation human haplotype map of over 3.1 million SNPs. Nature 449: 851–862.

Toro-Moyano, Isidro, Bienvenido Martinez-Navarro, Jordi Augusti, Caroline Souday, Jose Maria Bermudez de Castro, Maria Martinon-Torres, Beatriz Fajardo, et al. 2013. The oldest human fossil in Europe, from Orce (Spain). Journal of Human Evolution 65: 1–9.

Tsatskin, Alexander, and Mina Weinstein-Evron. 1994. The Jamal cave is not empty: recent discoveries in the Mount Carmel caves, Israel. Paleorient 20, 2: 119–128.

Wall, Jeffrey D., Kirk E. Lohmueller, and Vincent Plagnol. 2009. Detecting ancient admixture and estimating demographic parameters in multiple human populations. Molecular Biology and Evolution 26(8): 1823–1827.

Wall, Jeffrey D., Melinda A. Yang, Flora Jay, Sung K. Kim, Eric Y. Durand, Laurie S. Stevison, Christopher Gignoux, et al. 2013. Higher levels of Neanderthal ancestry in East Asians than in Europeans. Genetics 194: 199–209.

Weinstein-Evron, Mina, Alexander Tsatskin, Stephen Weiner, Ruth Shahack-Gross, Amos Frumkin, Reuven Yeshurun, and Yossi Zaidner. A Window into Early Middle Paleolithic Human Occupational Layers: Misliya Cave, Mount Carmel, Israel. 2012. PaleoAntropology: 202–228.

Yang, Melinda A., Anna-Sapfo Lalaspinas, Eric Y. Durand, and Montgomery Slatkin. 2012. Ancient Structure in Africa Unlikely to Explain Neandertal and Non-African Genetic Similarity. Molecular Biology and Evolution 29, 10, 2987–2995.

Zaidner, Yossi, Reuven Yeshuran, and Carolina Mallol. 2010. Early Pleistocene Hominins Outside of Africa: Recent Excavations at Bizat Ruhama, Israel. PaleoAnthropology 2010: 162–195.

Zaidner, Yossi. 2013. Adaptive Flexibility of Oldowan Hominins: Secondary Use of Flakes at Bizat Ruhama, Israel. PLOS ONE 8,6: 1–16.

